# Kinetic modeling of continuous meta-fermentation quantifies metabolic activity in a complex microbial system

**DOI:** 10.1101/2025.11.13.688382

**Authors:** Tomonori Koga, Kanta Kajimoto, Mitsuoki Ishizu, Hiroyuki Hamada, Hirokuni Miyamoto, Mugihito Oshiro, Kenji Sakai, Yukihiro Tashiro

**Author notes:** Corresponding author: Laboratory of Soil and Environmental Microbiology, Division of Systems Bioengineering, Department of Bioscience and Biotechnology, Faculty of Agriculture, Graduate School of Bioresources and Bioenvironmental Sciences, Kyushu University, Fukuoka 819-0395, Japan. E-mail address (Y. Tashiro).

## Abstract

One of the serious drawbacks of a complex microbial system is the difficulty in quantifying the metabolic activity of each microorganism. A kinetic model of predominant microbial species was constructed for continuous meta-fermentation in a complex microbial system at several dilution rates (D). The introduction of biomass and lactic acid inhibition terms improved model accuracies at D = 0.05 h^-1^ and 0.4 h^-1^, respectively. The coefficient of determination (R^2^) and root mean square error (RMSE) improved from 0.577 and 5.21 to 0.972 and 0.759, respectively, with the inhibition term at D = 0.05 h^-1^. The inhibition terms resulted in good R² (0.996) and RMSE (1.27) values at D = 0.4 h^-1^. By solving the Michaelis–Menten equation in the constructed models, the species flux (SF) was calculated to estimate the metabolic activity (formation and consumption) of each microorganism. At D = 0.05 h^-1^, *Caldibacillus hisashii* contributed to lactic acid production at 0.333 g/L/h, whereas *Clostridium cochlearium* consumed lactic acid at 0.203 g/L/h, suggesting cross-feeding of a metabolite. It is therefore possible to account for consumption that cannot be considered in gene-derived calculations, indicating that this is a promising analytical method for investigating the dynamic behavior of complex microbial systems. Kinetic models for continuous meta-fermentation at several D values were developed. A new metabolic analysis method is proposed to estimate the activity of microbial species in a complex microbial system.

## 1. Introduction

Organic acids are widely-used in food, cosmetics, and pharmaceuticals, and their demand is increasing annually. Lactic acid is a raw material for bioplastics and a platform chemical for conversion into other substances (1). Butyric acid is used as an aromatic ingredient, and the raw material is converted to esters in food and cosmetics (2). Due to the positive environmental impact, microbial organic acid production has attracted attention as an alternative to petrochemical production. Meta-fermentation, a fermentation process using complex microbial culture systems, has the advantages of utilization of multiple substrates, low cost, and no need for sterilization due to the low risk of microbial contamination, compared with a pure microbial system (3,4). However, complex microbial culture systems present challenges in elucidating the methods of microbial dynamics and lack technical knowledge for control.

Interactions among microorganisms in the environment occur through symbiosis, competition, and interdependence, which enables the decomposition and synthesis of substances (5,6). In a complex microbial culture system, the metabolic networks among microorganisms are more complex than those in a pure microbial system, resulting in a poor understanding of their interactions. Although several approaches, including metagenomics (7) and metatranscriptomics (8), have been developed for metabolic analysis in a complex microbial culture system, the obtained data (abundance of functional genomes and their expression levels) do not correspond to the actual metabolic activities of the targeted pathways of each microorganism (9). A thorough analysis of the metabolic activities of each microorganism during meta-fermentation is essential for efficient substance production and development of effective control strategies. Species-specific productivity (SSP) has been proposed to predict species level productivity using functional gene analysis based on the relative abundance of related genes (10). However, the SSP parameter only calculates the net productivity of the targeted pathways by each microorganism and cannot differentiate the formation flux of metabolites from their consumption flux, which would result in the underestimation or overestimation of actual productivity. Therefore, a new method should be established to quantify the metabolic activity of each microorganism by considering both formation and consumption fluxes.

Computational approaches, including genome-scale metabolic models and flux balance analysis (FBA), which stoichiometrically represent biochemical reactions within metabolic networks, have attracted attention as methods for elucidating these interactions (11–13). A kinetic model is an effective metabolic model that considers not only the stoichiometry of the metabolic network, but also its biological mechanisms, such as substrate and product inhibition. The kinetic model enables temporal and quantitative analysis of microbial metabolism and interactions with competition and symbiosis. The bacterial community structure changes depending on metabolite concentrations during cultivation (14), indicating the importance of dynamic and species-specific metabolic analyses.

A kinetic model using Monod’s equation has been reported for ethanol fermentation in a complex microbial system consisting of two species, *Scheffersomyces stipitis* and *Saccharomyces cerevisiae* (15). The constructed model showed good agreement with the experimental data for the substrates (glucose and xylose) and product (ethanol). A kinetic model should quantify the metabolic activity of each microorganism by considering both the formation and consumption fluxes in a complex microbial system. The current study aimed to construct a kinetic model of continuous meta-fermentation performance at several dilution rates dynamically and species-specifically and to develop a novel metabolic analysis method to predict productivity at the species level.

## 2. Material & Methods

### 2.1 Experimental data

The experimental data were obtained from a previous study (9). Briefly, a continuous meta-fermentation was conducted using 60 g/L glucose as a substrate and compost as a bacterial seed at a pH of 7.0 and 50 °C. Dilution rates (D) were changed stepwise from 0.4 h^-1^ to 0.05 h^-1^. Organic acids were quantified using high-performance liquid chromatography (HPLC) (Organic Acid Analyzer; Shimadzu, Kyoto, Japan) and glucose was determined using a biosensor (BF-7, Oji Scientific Instrument, Hyogo, Japan). Bacterial community structure was analyzed using 16S rRNA amplicon analysis on a MiSeq platform.

### 2.2 Kinetic modeling

Modeling and simulation were performed using WinBEST-KIT (Biochemical Engineering System analyzing Tool-KIT; Windows version) (16,17) which can construct and analyze reaction schemes expressed in terms of approximate rate functions for mass balance and enzymatic kinetics. Calculations of simultaneous differential equations were performed using the Gear method (18). WinBEST-KIT can estimate unknown kinetic parameters based on experimentally-observed time-course data (16,17), be used to construct metabolic models in pure culture systems, and identify key pathways for improving productivity (19,20) Information on metabolic pathways with predominant strains including *Caldibacillus hisashii* at D = 0.05 h^-1^ and 0.4 h^-1^, *Clostridium cochlearium* at D = 0.05 h^-1^, and *Heyndrickxia coagulans* at D = 0.4 h^-1^, were obtained from Kyoto Encyclopedia of Genes and Genomes (KEGG)(21). Metabolic pathways for the production of lactic acid, formic acid, acetic acid, and butyric acid from glucose were constructed for each of the three bacterial species. Metabolic reactions were conducted to simplify the analysis (Figs S1, 2). A reaction rate equation was constructed using the Michaelis–Menten equation. A biomass inhibition equation at D = 0.05 h^-1^ (Table S1), and lactic acid and biomass inhibition equations at D = 0.4 h^-1^ (Table S3) were introduced. Organic acids were assumed to be produced and consumed by the respective species, and then shared between species (Tables S2, S4). The values of kinetic parameters in the model were estimated using parameter estimations to match experimental data at D = 0.05 h^-1^ and 0.4 h^-1^, respectively. The inhibition terms models parameters at D = 0.05 h^-1^ and 0.4 h^-1^ are shown in Table S5 and S6 respectively.

### 2.4 Validation of the models and calculation of species flux

The coefficient of determination (R²) and root mean square error (RMSE) between the simulation results and experimental data were calculated to validate the accuracy of the constructed models. R² was calculated as given in Equation (1):

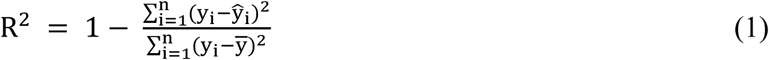

where n is the number of data points; y_i_ and ŷ_i_ are the values of the experimental and simulation data at the same fermentation time, respectively; and ȳ is the average value of the experimental data.

RMSE was calculated as given in Equation (2):

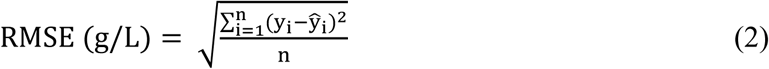

The Species flux (SF) was calculated by summing the reaction rate equations representing the formation and consumption rate of the target substance (Equation 3).

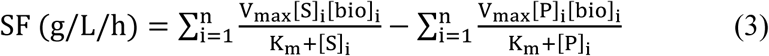

where [S] is substrate concentration, [bio] is the concentration of each microbial species, and [P] is the concentration of the target product.

The species-specific productivity (SSP) was calculated using (Eq. 4) (10):

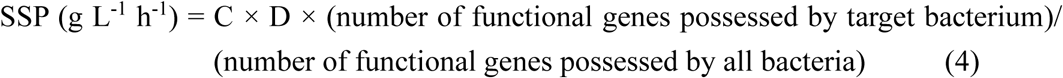

where C is the organic acid concentration (g L^-1^) and D is the dilution rate (h^-1^).

## 3. Results

### 3.1 Kinetic model at D = 0.05 h^-1^

In this study, two predominant species in continuous meta-fermentation were included at D = 0.05 h^-1^ as in a previous study (10), and a dynamic model capable of representing the fermentation performances of the substrate and products was constructed. At D = 0.05 h^-1^, *C. hisashii* (average, 60.6%) and *C. cochlearium* (average, 34.0%) were predominant species, with production of 2.69 g/L lactic acid, 6.96 g/L butyric acid, 3.95 g/L acetic acid, and 2.28 g/L formic acid and consumption of 45.3 g/L glucose (10). Closely-related species of *C. hisashii* are known to produce lactic acid as the main product, in addition to the byproducts of formic acid and acetic acid by heterolactic fermentation. *C. cochlearium* produced acetic acid and butyric acid (22). At D = 0.05 h^-1^, the R² and RMSE values of the model without a biomass inhibition term were: lactic acid, 0.730 and 1.10; formic acid, 0.524 and 0.399; acetic acid, 0.156 and 0.505; butyric acid, 0.906 and 0.589; glucose, -8.75 and 11.6; and overall, 0.577 and 5.21, respectively (Table S7). The R² value for glucose was negative, indicating a markedly poor prediction. In contrast, the model with a biomass inhibition term improved the R² and RMSE values of metabolites: 0.724 and 0.590 for lactic acid, 0.888 and 0.0333 for formic acid, 0.653 and 0.0555 for acetic acid, 0.911 and 0.456 for butyric acid, and 0.864 and 1.74 for glucose, indicating sufficient accuracy for each metabolite. As a result of improving the values of metabolites by introducing a biomass inhibition term, the overall R² increased to 0.972 from 0.577 and the RMSE decreased to 0.759 from 5.21. The simulation data from the model with the biomass inhibition term showed qualitative agreement with the experimental data over time. The biomass inhibition term could therefore be important in establishing a good kinetic model; biomass inhibition contributes to metabolic activities in continuous meta-fermentation at D = 0.05 h^-1^.

### 3.2 Kinetic model at D = 0.4 h^-1^

At D = 0.4 h^-1^, the relative abundance of *C. hisashii* and *H. coagulans* was 89.3% and 10.6%, respectively, with productions of 6.60 g/L lactic acid, 0.740 g/L acetic acid, and 0.560 g/L formic acid without butyric acid production, and consumption of 16.9 g/L glucose (10). *H. coagulans* produces lactic acid via homolactic fermentation (23). At D = 0.4 h^-1^, the R² and RMSE values without inhibition terms were: lactic acid, 0.668 and 0.326; formic acid, 0.171 and 0.145; and acetic acid, -0.119 and 0.188; glucose, 0.0345 and 44.7, respectively (Table S8). By introducing the terms biomass and lactic acid inhibition terms, the R² and RMSE values were found to be 0.795 and 0.293 for lactic acid, 0.0603 and 0.254 for formic acid, 0.0710 and 0.171 for acetic acid, and 0.251 and 2.50 for glucose, respectively (Table S8). The improvement in these values for each metabolite resulted in a much better overall R² and RMSE of 0.996 and 1.27, respectively, with the introduction of terms for biomass and lactic acid inhibition, than those without of 0.0363 and 22.4, respectively (Fig. 2; Table S8). Simulation data from the model with biomass and lactic acid inhibition terms showed qualitative agreement with the experimental data over time (Fig. 3b). These results strongly suggest that introducing inhibition kinetics would substantially enhance the validity of dynamic simulations of metabolic activities, and that biomass and lactic acid inhibition affect the metabolic activities of both species in continuous meta-fermentation at D = 0.4 h^-1^.

**Figure 1.**
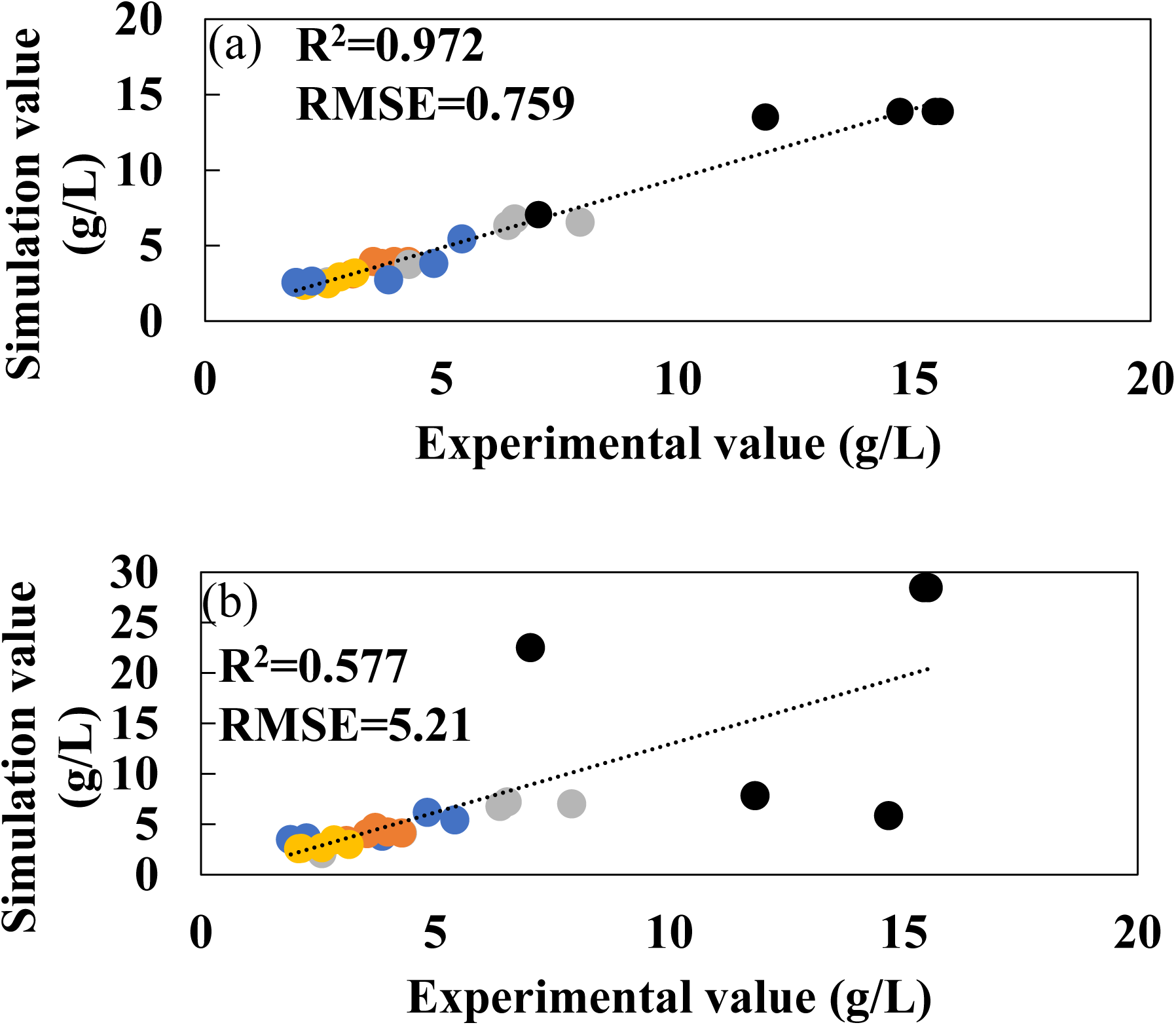
Experimental (•) and simulation (–) concentrations of lactic acid (blue), formic acid (yellow), acetic acid (orange), butyric acid (gray), and glucose (black) in continuous meta-fermentation at D of 0.05 h^-1^ (a) without inhibition terms (b) with inhibition terms.

**Figure 2.**
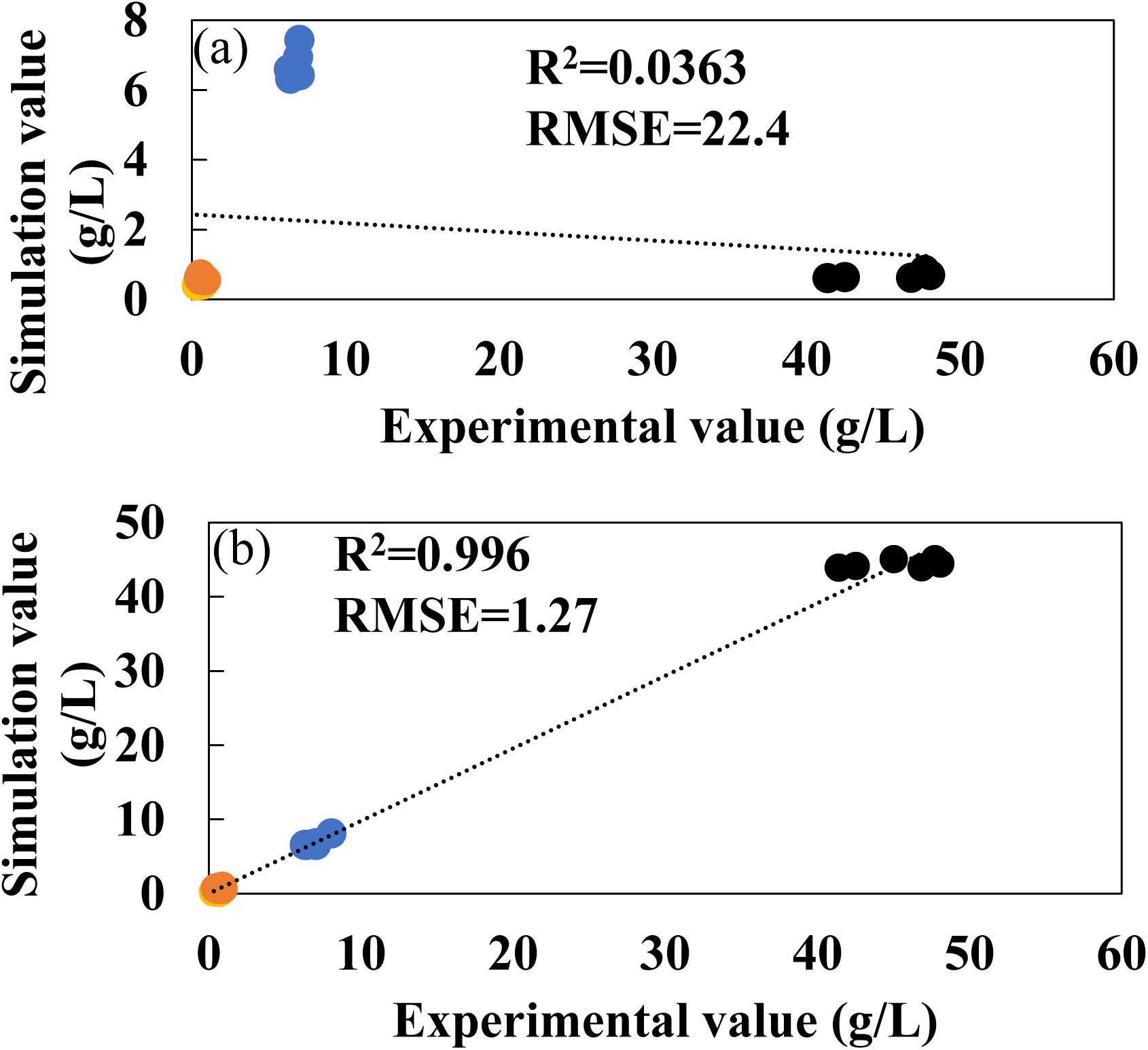
Experimental (•) and simulation (–) concentrations of lactic acid (blue), formic acid (yellow), acetic acid (orange), and glucose (black) in continuous meta-fermentation at D of 0.4 h^-1^ (a) without inhibition terms (b) with inhibition terms.

**Figure 3.**
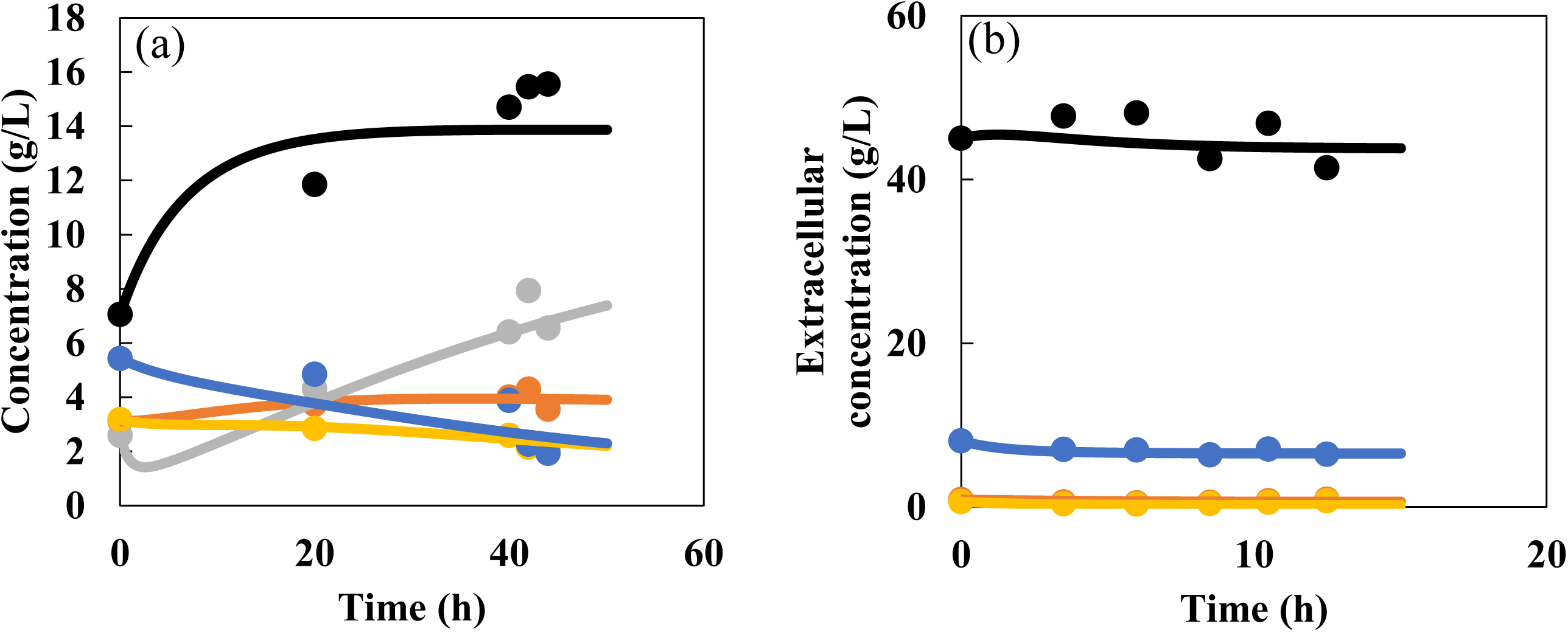
Time-course of experimental (•) and simulation (–) concentrations of inhibition terms lactic acid (blue), formic acid (yellow), acetic acid (orange), butyric acid (gray), and glucose (black) in continuous meta-fermentation at (a) D = 0.05 h^-1^ and (b) D = 0.4 h^-1^.

### 3.3 Quantification of species flux for products by constructed kinetic models

By solving Michaelis–Menten equations, the Species flux (SF) was calculated using Eq. (3). At D = 0.05 h^-1^, SF of *C. hisashii* for lactic acid, formic acid, and acetic acid productions was 0.333, 0.132, and 0.168 g/L/h, respectively (Fig. 4a). The SF of *C. cochlearium* for the production of acetic acid (0.0286 g/L/h) and butyric acid (0.333 g/L/h) was obtained, whereas the consumption of lactic acid and formic acid was observed at SF of 0.203 and 0.0124 g/L/h, respectively. Cross-feeding of lactic acid between *C. cochlearium* and *C. hisashii* occurred, and *C. cochlearium* contributed to butyric acid production by consuming the lactic acid produced by *C. hisashii* at D = 0.05 h^-1^.

**Figure 4.**
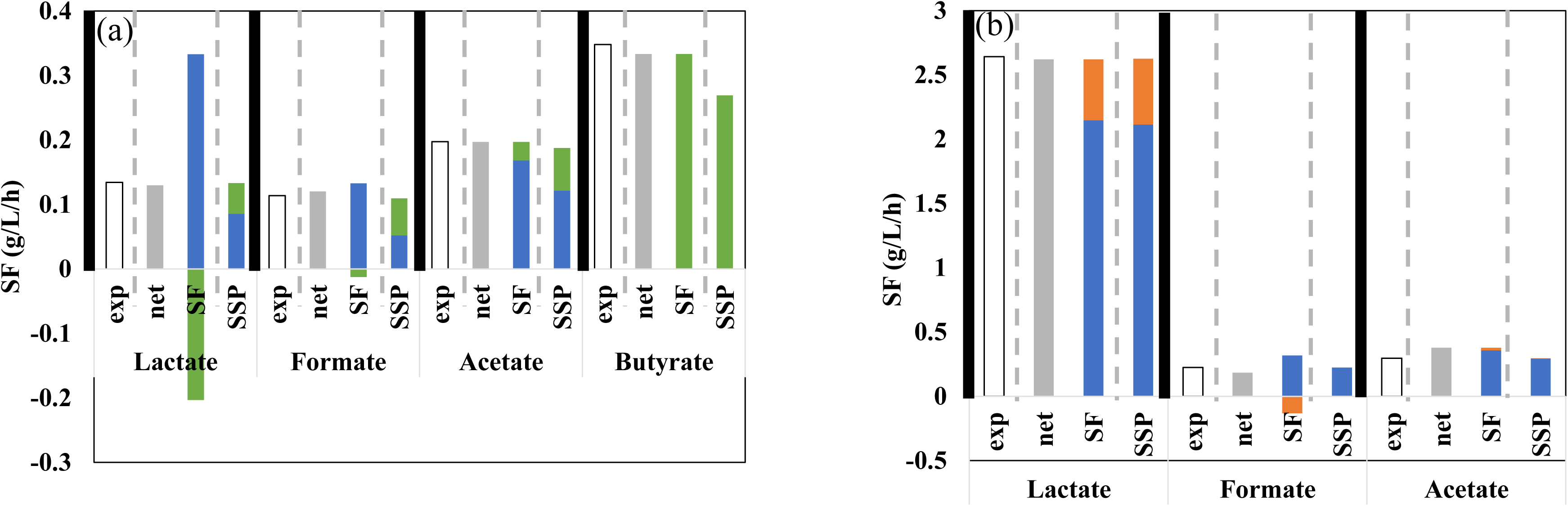
Experimental productivity (white), net productivity of SF (gray), SF, and SSP of each organic acid produced by *C. hisashii* (blue), *C. cochlearium* (green) and *H. coagulans* at (a) D = 0.05 h^-1^ and (b) D = 0.4 h^-1^.

Figure 4b shows the SF values for each organic acid at D = 0.4 h^-1^: *C. hisashii*, production for lactic acid (2.14 g/L/h), formic acid (0.317 g/L/h), and acetic acid (0.357 g/L/h); *H. coagulans*, production of lactic acid (0.472 g/L/h). The metabolic activities of *C. hisashii* for lactic acid, formic acid, and acetic acid production changed depending on D values between 0.05 h^-1^ and 0.4 h^-1^.

The SSP and SF values were compared (Fig. 4). At D = 0.4 h^-1^, similar SSP and SF values of *C. hisashii* and *H. coagulans* were obtained for production of lactic acid, formic acid, and acetic acid. Thus, SSP can be used to quantify the metabolic activity of each microorganism under conditions without the consumption of metabolites. The values of *C. cochlearium* and *C. hisashii* for lactic acid, formic acid, acetic acid, and butyric acid differed between SSP and SF. The behavior of lactic acid was drastically different. Metabolite consumption is not considered in the calculation of SSP in Eq. (4), which would lead to an incorrect and incalculable parameter under conditions of metabolite cross-feeding among microorganisms. The metabolic activity of each microorganism was estimated by modifying parameters such as the dilution rate in the kinetic model without experiments.

Consequently, the constructed model enabled the dynamic and quantitative estimation of metabolic activity at the species level during meta-fermentation.

## 4. Discussion

A method to quantify the metabolic activities of production and consumption with each microbial species in complex microbial systems was developed, particularly that of meta-fermentation for organic acid production. The estimation of metabolic activities at the microbial species level is important for the optimization of bioprocesses and the development of control strategies in complex microbial systems as well as pure microbial systems (20,24). However, there have been few studies on complex microbial systems, and no theory has been developed for the optimization and control of meta-fermentation. The current study attempted to quantify the metabolic activities of predominant microbial species by constructing a kinetic model and proposes a new approach for complex microbial systems.

The kinetic models developed in this study could simulate the metabolites (one substrate, four products) over time and quantitatively in continuous meta-fermentation at D = 0.05 h^-1^ and 0.4 h^-1^. The overall R^2^ and RMSE values for this model of D = 0.05 h^-1^ and 0.4 h^-1^ were 0.972 and 0.759, and 0.996 and 1.27, respectively (Figs 1, 3a; Table S7). Table 1 shows a comparison of the metabolic models for complex and pure microbial systems. Most studies have targeted pure culture systems, including fermentative products such as ethanol, butanol, and lactic acid. For example, correlation coefficients of R^2^ = 0.917–0.926 were obtained using a kinetic model of butanol production from glucose or mixed substrates (glucose and xylose) in pure culture systems with *Clostridium beijerinckii* (25). To the best of our knowledge, highly accurate metabolic models have not been developed for complex microbial systems. This study successfully developed a model with accuracy equivalent to that of pure culture systems.(15,19,20,25–28)

**Table 1.**
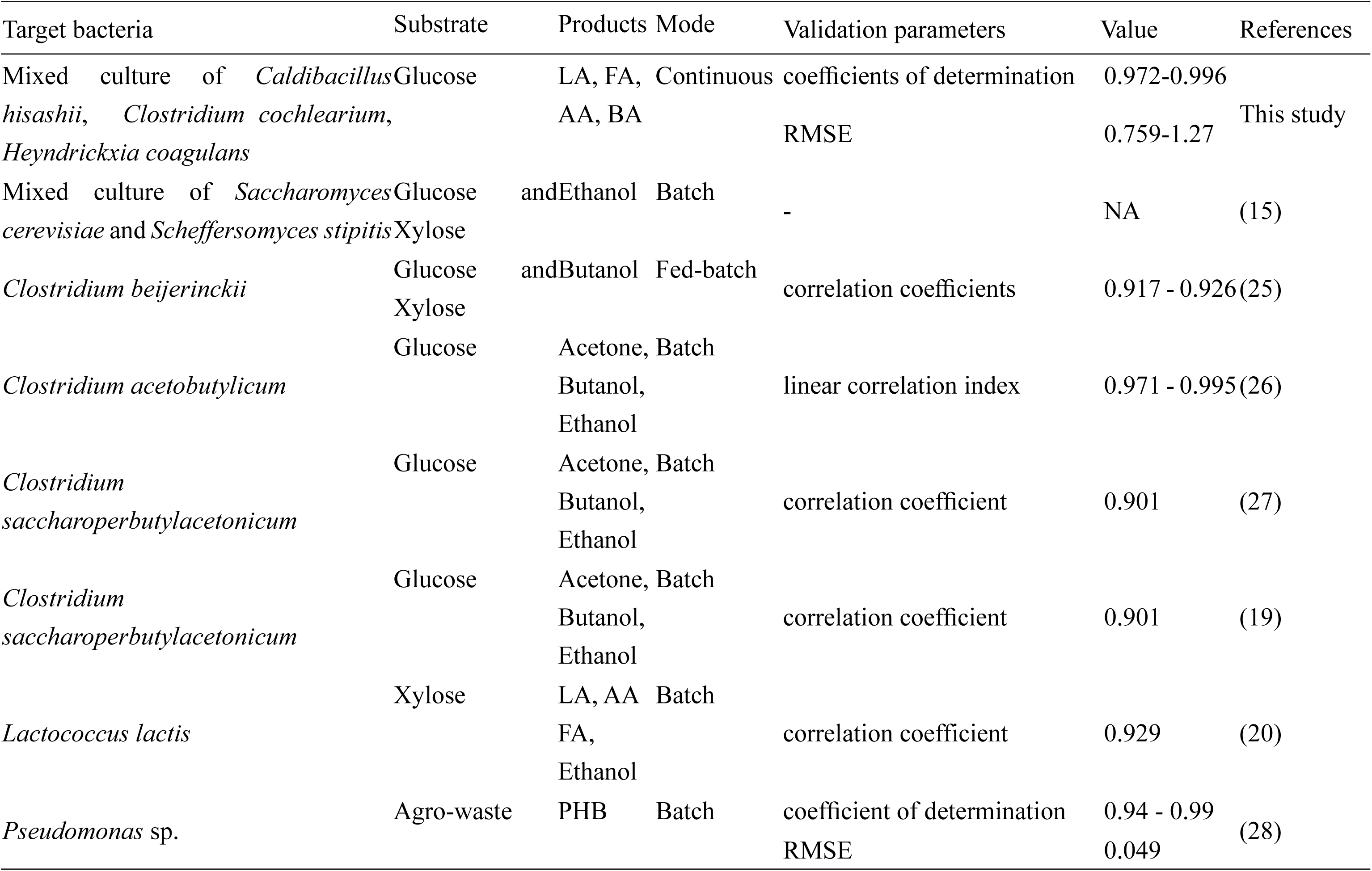
Modeling of various fermentation production and validation in literature.

The model presented in this study introduced the non-competitive inhibition terms with lactic acid and biomass inhibition, which led to improvements in the overall R^2^ and RMSE values from 0.577 and 5.21 to 0.972 and 0.759 at D = 0.05 h^-1^ and 0.0363 and 22.4 to 0.996 and 1.27 at D = 0.4 h^-1^, respectively (Table S7, 8). In previous studies on pure culture systems, metabolic models with various modifications were reported to be more accurate (29,30). In particular, the non-competitive Michaelis–Menten equation has often been applied because lactic acid and affect enzymatic activity via non-competitive regulatory mechanisms in microbial cells (31). In addition, a biomass inhibition term is commonly incorporated as an empirical means to limit excessive biomass accumulation under conditions of continuous substrate feeding (32). Thus, these results suggest that lactic acid noncompetitively inhibit metabolic activities in this complex microbial system.

Cross-feeding, the exchange of metabolites, such as energy and nutrients, among different microbial species, has generally been reported in complex microbial systems (33). At D = 0.05 h^-1^, the lactic acid produced by *C. hisashii* with SF of 0.333 g/L/h was consumed by *C. cochlearium* with SF of 0.203 g/L/h (Figs 4a). The constructed kinetic model provided temporal and quantitative estimations of the formation and consumption of lactic acid by each microbial species (Figs S3, S4), which led to the hypothesis of cross-feeding during continuous meta-fermentation. These results demonstrate the potential of this new method for analyzing complex metabolic interactions, such as competition and symbiosis, in complex microbial systems.

Although many studies have analyzed metabolism in pure culture systems using ^13^C-metabolic flux analysis (MFA) (34,35), ^13^C-MFA is not suitable for complex microbial systems because there is no method for analyzing the consumption of substrates and products at the microbial species level. Metagenomic and metatranscriptomic analyses have been used for genomic analysis. However, it is difficult to quantify the dynamics and metabolic activities of microbial species. This is due to differences in the genome, transcription levels, and enzymatic activities (9). A new method is proposed for quantifying the metabolic activity of microbial species over time as a parameter of SF. This method enables the estimation of substance-mediated interactions between microorganisms and allows the numerical calculation of metabolic fluxes, even for complex microbial systems involving three or more microbial species.

Although the field of microbial engineering using pure microbial systems has been systematized, complex microbial engineering lacks sufficient theoretical and technical knowledge (10). Although this study established a new method for analyzing the metabolic activity and function of each microbial species in a complex system, further research is needed to apply this approach to optimize substance production and control microbial community structure during meta-fermentation.

Kinetic models were constructed to represent the behavior of continuous meta-fermentation at several dilution rates, and the species flux (SF) was calculated. The temporal and quantitative production of organic acids by *C. hisashii* and the consumption of lactic acid by *C. cochlearium* were hypothesized. This study suggests that this new metabolic analysis method can be used to calculate species-level productivity, while considering consumption in complex microbial systems involving meta-fermentation.

## Acknowledgments

This work was partly supported by JSPS KAKENHI Grant-in-Aid for Scientific Research (B) Grant Number JP 23H03595, JST SPRING Grant Number JPMJSP2136.

## Author contributions

Tomonori Koga: Original Draft Preparation, Resources, Data Curation Formal Analysis, Investigation, Visualization. Kanta Kajimoto: Methodology, Software, Validation, Formal Analysis, Investigation, Visualization. Mitsuoki Ishizu: Methodology, Software, Validation, Formal Analysis, Investigation. Yukihiro Tashiro Conceptualization, Review & Editing, Funding Acquisition, Project Administration, Supervision. Kenji Sakai: Supervision. Hiroyuki Hamada: Supervision. Mugihito Oshiro: Supervision.

