## Supplementary material for "Kinetic modeling of continuous meta-fermentation quantifies metabolic activity in a complex microbial system": suppleTable

RIKEN IMS, Kanagawa 230-0045, Japan,[3]

Graduate School of Horticulture, Chiba University, Chiba 271-8510, Japan,[4]

Sermas Co., Ltd., Chiba 272-0015, Japan,[5]

Japan Eco-science (Nikkan Kagaku) Co. Ltd., Chiba 263-8522, Japan,[6]

and Laboratory of Microbial Environmental Protection, Tropical Microbiology Unit, Center for International Education and Research of Agriculture, Faculty of Agriculture, Kyushu University, Fukuoka 819-0395, Japan,[7]

Table S1. The rate equations of each metabolic reaction at D = 0.05 $h^{-1}$.

| | |
|---|---|
| $r_{1\_A} = \frac{V_{max1\_A}[Glucose][Biomass\_A]}{([Glucose] + K_{m1\_A})(1 + \frac{[Biomass\_A]}{K_{iL\_A}})}$ | $r_{1\_C} = \frac{V_{max1\_C}[Glucose][Biomass\_C]}{([Glucose] + K_{m1\_C})(1 + \frac{[Biomass\_C]}{K_{iL\_C}})}$ |
| $r_{2\_A} = \frac{V_{max2\_A}[G3P\_A][Biomass\_A]}{K_{m2\_A} + [G3P\_A]}$ | $r_{2\_C} = \frac{V_{max2\_C}[G3P\_C][Biomass\_C]}{K_{m2\_C} + [G3P\_C]}$ |
| $r_{3\_A} = \frac{V_{max3\_A}[LA][Biomass\_A]}{K_{m3\_A} + [LA]}$ | $r_{3\_C} = \frac{V_{max3\_C}[LA][Biomass\_C]}{K_{m3\_C} + [LA]}$ |
| $r_{4\_A} = \frac{V_{max4\_A}[Pyruvate\_A][Biomass\_A]}{K_{m4\_A} + [Pyruvate\_A]}$ | $r_{4\_C} = \frac{V_{max4\_C}[Pyruvate\_C][Biomass\_C]}{K_{m4\_C} + [Pyruvate\_C]}$ |
| $r_{5\_A} = \frac{V_{max5\_A}[Pyruvate\_A][Biomass\_A]}{K_{m5\_A} + [Pyruvate\_A]}$ | $r_{5\_C} = \frac{V_{max5\_C}[Pyruvate\_C][Biomass\_C]}{K_{m5\_C} + [Pyruvate\_C]}$ |
| $r_{6\_A} = \frac{V_{max6\_A}[Pyruvate\_A][Biomass\_A]}{K_{m6\_A} + [Pyruvate\_A]}$ | $r_{6\_C} = \frac{V_{max6\_C}[Pyruvate\_C][Biomass\_C]}{K_{m6\_C} + [Pyruvate\_C]}$ |
| $r_{7\_A} = \frac{V_{max7\_A}[AA][Biomass\_A]}{K_{m7\_A} + [AA]}$ | $r_{7\_C} = \frac{V_{max7\_C}[AA][Biomass\_C]}{K_{m7\_C} + [AA]}$ |
| $r_{8\_A} = \frac{V_{max8\_A}[ACoA\_A][Biomass\_A]}{K_{m8\_A} + [ACoA\_A]}$ | $r_{8\_C} = \frac{V_{max8\_C}[ACoA\_C][Biomass\_C]}{K_{m8\_C} + [ACoA\_C]}$ |
| $r_{9\_A} = \frac{V_{max9\_A}[ACoA\_A][Biomass\_A]}{K_{m9\_A} + [ACoA\_A]}$ | $r_{9\_C} = \frac{V_{max9\_C}[ACoA\_C][Biomass\_C]}{K_{m9\_C} + [ACoA\_C]}$ |
| $r_{m\_A} = \frac{V_{maxm\_A}[FA][Biomass\_A]}{K_{mm\_A} + [FA]}$ | $r_{m\_C} = \frac{V_{maxm\_C}[FA][Biomass\_C]}{K_{mm\_C} + [FA]}$ |
| $r_{10\_C} = \frac{V_{max10\_C}[ACoA\_C][Biomass\_C]}{K_{m10\_C} + [ACoA\_C]}$ | $r_{11\_C} = \frac{V_{max11\_C}[BA][Biomass\_C]}{K_{m11\_C} + [BA]}$ |
| $r_{12\_C} = \frac{V_{max12\_C}[BCoA\_C][Biomass\_C]}{K_{m12\_C} + [BCoA\_C]}$ | $outflow_1 = D[Glucose]$ |
| $outflow_2 = D[Lactate]$ | $outflow_3 = D[Formate]$ |
| $outflow_4 = D[Acetate]$ | $outflow_5 = D[Butyrate]$ |
| $inflow = D[Stoc\_Glucose]$ | |

Table S2. The reaction balance of the target metabolites at D = 0.05 $h^{-1}$.

| | |
|---|---|
| $\frac{d[Glucose]}{dt} = Inflow - Outflow_1 - r_{1\_A} - r_{1\_C}$ | $\frac{d[Lactate]}{dt} = -Outflow_2 - r_{3\_A} - r_{3\_C} + r_{4\_A} + r_{4\_c}$ |
| $\frac{d[Formate]}{dt} = -Outflow_3 + r_{5\_A} + r_{5\_C} - r_{m\_A} - r_{m\_C}$ | $\frac{d[Acetate]}{dt} = -Outflow_4 - r_{7\_A} - r_{7\_C} + r_{8\_A} + r_{8\_C}$ |
| $\frac{d[G3P\_A]}{dt} = r_{1\_A} - r_{2\_A}$ | $\frac{d[G3P\_C]}{dt} = r_{1\_C} - r_{2\_C}$ |
| $\frac{d[Pyruvate\_A]}{dt} = r_{2\_A} + r_{3\_A} - r_{4\_A} - r_{5\_A} - r_{6\_A}$ | $\frac{d[Pyruvate\_C]}{dt} = r_{2\_C} + r_{3\_C} - r_{4\_C} - r_{5\_C} - r_{6\_C}$ |
| $\frac{d[ACoA\_A]}{dt} = r_{5\_A} + r_{6\_A} + r_{7\_A} - r_{8\_A} - r_{9\_A}$ | $\frac{d[ACoA\_C]}{dt} = r_{5\_C} + r_{6\_C} + r_{7\_C} - r_{8\_C} - r_{9\_C}$ |
| $\frac{d[Biomass\_A]}{dt} = -Death - Outflow_5 + r_{9\_A}$ | $\frac{d[Biomass\_C]}{dt} = -Death - Outflow_5 + r_{9\_C}$ |
| $\frac{d[BCoA\_C]}{dt} = r_{10\_C} + r_{11\_C} - r_{12\_C}$ | $\frac{d[Butyrate]}{dt} = -Outflow_6 - r_{11\_C} + r_{12\_C}$ |

Table S3. The rate equations of each metabolic reaction at D = 0.4 h[-1].

| | |
|---|---|
| $r_{1\_A} = \frac{V_{max1\_A}[\text{Glucose}][\text{Biomass\_A}]}{([\text{Glucose}] + K_{m1\_A})(1 + \frac{[\text{Biomass\_A}]}{K_{iB\_A}})(1 + \frac{[\text{LA}]}{K_{iL\_A}})}$ | $r_{1\_W} = \frac{V_{max1\_H}[\text{Glucose}][\text{Biomass\_W}]}{([\text{Glucose}] + K_{m1\_W})(1 + \frac{[\text{Biomass\_W}]}{K_{iB\_W}})(1 + \frac{[\text{LA}]}{K_{iL\_W}})}$ |
| $r_{2\_A} = \frac{V_{max2\_A}[\text{G3P\_A}][\text{Biomass\_A}]}{K_{m2\_A} + [\text{G3P\_A}]}$ | $r_{2\_W} = \frac{V_{max2\_W}[\text{G3P\_W}][\text{Biomass\_W}]}{K_{m2\_W} + [\text{G3P\_W}]}$ |
| $r_{3\_A} = \frac{V_{max3\_A}[\text{LA}][\text{Biomass\_A}]}{K_{m3\_A} + [\text{LA}]}$ | $r_{3\_W} = \frac{V_{max3\_W}[\text{LA}][\text{Biomass\_W}]}{K_{m3\_W} + [\text{LA}]}$ |
| $r_{4\_A} = \frac{V_{max4\_A}[\text{Pyruvate\_A}][\text{Biomass\_A}]}{K_{m4\_A} + [\text{Pyruvate\_A}]}$ | $r_{4\_W} = \frac{V_{max4\_W}[\text{Pyruvate\_W}][\text{Biomass\_W}]}{K_{m4\_W} + [\text{Pyruvate\_W}]}$ |
| $r_{5\_A} = \frac{V_{max5\_A}[\text{Pyruvate\_A}][\text{Biomass\_A}]}{K_{m5\_A} + [\text{Pyruvate\_A}]}$ | $r_{5\_W} = \frac{V_{max5\_W}[\text{Pyruvate\_W}][\text{Biomass\_W}]}{K_{m5\_W} + [\text{Pyruvate\_W}]}$ |
| $r_{6\_A} = \frac{V_{max6\_A}[\text{Pyruvate\_A}][\text{Biomass\_A}]}{K_{m6\_A} + [\text{Pyruvate\_A}]}$ | $r_{6\_W} = \frac{V_{max6\_W}[\text{Pyruvate\_W}][\text{Biomass\_W}]}{K_{m6\_W} + [\text{Pyruvate\_W}]}$ |
| $r_{7\_A} = \frac{V_{max7\_A}[\text{AA}][\text{Biomass\_A}]}{K_{m7\_A} + [\text{AA}]}$ | $r_{7\_W} = \frac{V_{max7\_W}[\text{AA}][\text{Biomass\_W}]}{K_{m7\_W} + [\text{AA}]}$ |
| $r_{8\_A} = \frac{V_{max8\_A}[\text{ACoA\_A}][\text{Biomass\_A}]}{K_{m8\_A} + [\text{ACoA\_A}]}$ | $r_{8\_W} = \frac{V_{max8\_W}[\text{ACoA\_W}][\text{Biomass\_W}]}{K_{m8\_W} + [\text{ACoA\_W}]}$ |
| $r_{9\_A} = \frac{V_{max9\_A}[\text{ACoA\_A}][\text{Biomass\_A}]}{K_{m9\_A} + [\text{ACoA\_A}]}$ | $r_{9\_W} = \frac{V_{max9\_W}[\text{ACoA\_W}][\text{Biomass\_W}]}{K_{m9\_W} + [\text{ACoA\_W}]}$ |
| $r_{m\_A} = \frac{V_{maxm\_A}[\text{FA}][\text{Biomass\_A}]}{K_{mm\_A} + [\text{FA}]}$ | $r_{m\_W} = \frac{V_{maxm\_W}[\text{FA}][\text{Biomass\_W}]}{K_{mm\_W} + [\text{FA}]}$ |
| $\text{outflow}_1 = D[\text{Glucose}]$ | $\text{outflow}_2 = D[\text{Lactate}]$ |
| $\text{outflow}_3 = D[\text{Formate}]$ | $\text{outflow}_4 = D[\text{Acetate}]$ |
| $\text{inflow} = D[\text{Stoc\_Glucose}]$ | |

Table S4. The reaction balance of the target metabolites at D = 0.4 $h^{-1}$.

| | |
|---|---|
| $\frac{d[Glucose]}{dt} = Inflow - Outflow_1 - r_{1\_A} - r_{1\_W}$ | $\frac{d[Lactate]}{dt} = -Outflow_2 - r_{3\_A} - r_{3\_W} + r_{4\_A} + r_{4\_W}$ |
| $\frac{d[Formate]}{dt} = -Outflow_3 + r_{5\_A} + r_{5\_W} - r_{m\_A} - r_{m\_W}$ | $\frac{d[Acetate]}{dt} = -Outflow_4 - r_{7\_A} - r_{7\_W} + r_{8\_A} + r_{8\_W}$ |
| $\frac{d[G3P\_A]}{dt} = r_{1\_A} - r_{2\_A}$ | $\frac{d[G3P\_W]}{dt} = r_{1\_W} - r_{2\_W}$ |
| $\frac{d[Pyruvate\_A]}{dt} = r_{2\_A} + r_{3\_A} - r_{4\_A} - r_{5\_A} - r_{6\_A}$ | $\frac{d[Pyruvate\_W]}{dt} = r_{2\_W} + r_{3\_W} - r_{4\_W} - r_{5\_W} - r_{6\_W}$ |
| $\frac{d[ACoA\_A]}{dt} = r_{5\_A} + r_{6\_A} + r_{7\_A} - r_{8\_A} - r_{9\_A}$ | $\frac{d[ACoA\_W]}{dt} = r_{5\_W} + r_{6\_W} + r_{7\_W} - r_{8\_W} - r_{9\_W}$ |
| $\frac{d[Biomass\_A]}{dt} = -Death - Outflow_5 + r_{9\_A}$ | $\frac{d[Biomass\_W]}{dt} = -Death - Outflow_5 + r_{9\_W}$ |

Table S5 Kinetic parameters estimated in the kinetic models for *Caldibacillus hisashii* and *Clostridium cochlearium* at D = 0.4 $h^{-1}$

| Symbol Name | *C. cochlearium* | | | *C. hisashii* | | |
|---|---|---|---|---|---|---|
| | Va | Km | KiL | Va | Km | KiL |
| R1 | 7 | 80 | 1.5 | 9.5 | 60 | 1.5 |
| R2 | 10 | 1 | | 10 | 1 | |
| R3 | 0.0752 | 1 | | 0.01 | 0.01 | |
| R4 | 1.8 | 10 | | 2.28 | 0.8 | |
| R5 | 7 | 1 | | 4.17 | 1.06 | |
| R6 | 21 | 1 | | 10 | 3 | |
| R7 | 0.071 | 59 | | 0.0683 | 50 | |
| R8 | 4.04 | 2.2 | | 11.5 | 2.2 | |
| R9 | 3.01 | 1 | | 4.3 | 1 | |
| R10 | 12.4 | 2 | | - | - | |
| R11 | 1.22 | 20 | | - | - | |
| R12 | 1.06 | 12 | | - | - | |
| Metab | 0.096 | 1.1 | | 0.056 | 1.53 | |

Table S6 Kinetic Parameters estimated in the kinetic models for *Caldibacillus hisashii* and *Heyndrickxia coagulans* at D = 0.4 $h^{-1}$

| Symbol | *H. coagulans* | | | | *C. hisashii* | | | |
|---|---|---|---|---|---|---|---|---|
| Name | Va | Km | KiB | KiL | Va | Km | KiB | KiL |
| R1 | 480 | 2.92 | 1 | 1 | 2280 | 12.6 | 1.026 | 0.9919 |
| R2 | 173.8 | 1 | | | 193.4 | 1 | | |
| R3 | 0.086 | 8.3 | | | 0.121 | 100 | | |
| R4 | 38.4 | 0.789 | | | 77.5 | 1.004 | | |
| R5 | 12.5 | 100 | | | 24.192 | 2.4 | | |
| R6 | 48 | 0.99 | | | 250 | 3 | | |
| R7 | 0.1 | 90 | | | 0.1 | 15 | | |
| R8 | 3 | 15 | | | 7.96 | 0.92 | | |
| R9 | 99.9 | 34.41 | | | 120 | 3.78 | | |
| Metab | 0.0767 | 0.982 | | | 0.01021 | 13.95 | | |

Table S7 Correlation of determination between inhibition equation models and non-inhibition equation models for each substance with $D = 0.05\ h^{-1}$

| Metabolites | $R^2$ | | RMSE (g/L) | |
|---|---|---|---|---|
| | Inhibited model | non inhibited model | Inhibited model | non inhibited model |
| LA | 0.724 | 0. 730 | 0.590 | 1.10 |
| FA | 0.888 | 0.524 | 0.0333 | 0.399 |
| AA | 0.653 | 0.156 | 0.0555 | 0.505 |
| BA | 0.911 | 0.906 | 0.456 | 0.589 |
| Glc | 0.864 | -8.75 | 1.74 | 11.6 |
| All | 0.972 | 0.577 | 0.759 | 5.21 |

Table S8 Correlation coefficients between inhibition equation models and non-inhibition equation models for each substance with D = 0.4 $h^{-1}$

| Metabolites | $R^2$ | | RMSE (g/L) | |
|---|---|---|---|---|
| | Inhibited model | non inhibition model | Inhibited model | non inhibited model |
| LA | 0.795 | 0.668 | 0.293 | 0.326 |
| FA | 0.0603 | 0.171 | 0.254 | 0.145 |
| AA | 0.0710 | -0.119 | 0.171 | 0.188 |
| Glc | 0.251 | 0.0345 | 2.50 | 44.7 |
| All | 0.996 | 0.0363 | 1.27 | 22.4 |
