## Supplementary material for "Kinetic modeling of continuous meta-fermentation quantifies metabolic activity in a complex microbial system": suppleFig

Affiliations
Laboratory of Soil and Environmental Microbiology, Division of Systems Bioengineering, Department of Bioscience and Biotechnology, Faculty of Agriculture, Graduate School of Bioresources and Bioenvironmental Sciences, Kyushu University, Fukuoka 819-0395, Japan,[1]
RIKEN IMS, Kanagawa 230-0045, Japan,[2]
Graduate School of Horticulture, Chiba University, Chiba 271-8510, Japan,[3]
Sermas Co., Ltd., Chiba 272-0015, Japan,[4]
Japan Eco-science (Nikkan Kagaku) Co. Ltd., Chiba 263-8522, Japan,[5]
and Laboratory of Microbial Environmental Protection, Tropical Microbiology Unit, Center for International Education and Research of Agriculture, Faculty of Agriculture, Kyushu University, Fukuoka 819-0395, Japan,[6]

[*] Corresponding author:
Laboratory of Soil and Environmental Microbiology,
Division of Systems Bioengineering, Department of Bioscience and Biotechnology,
Faculty of Agriculture, Graduate School of Bioresources and Bioenvironmental Sciences, Kyushu University, Fukuoka 819-0395, Japan.

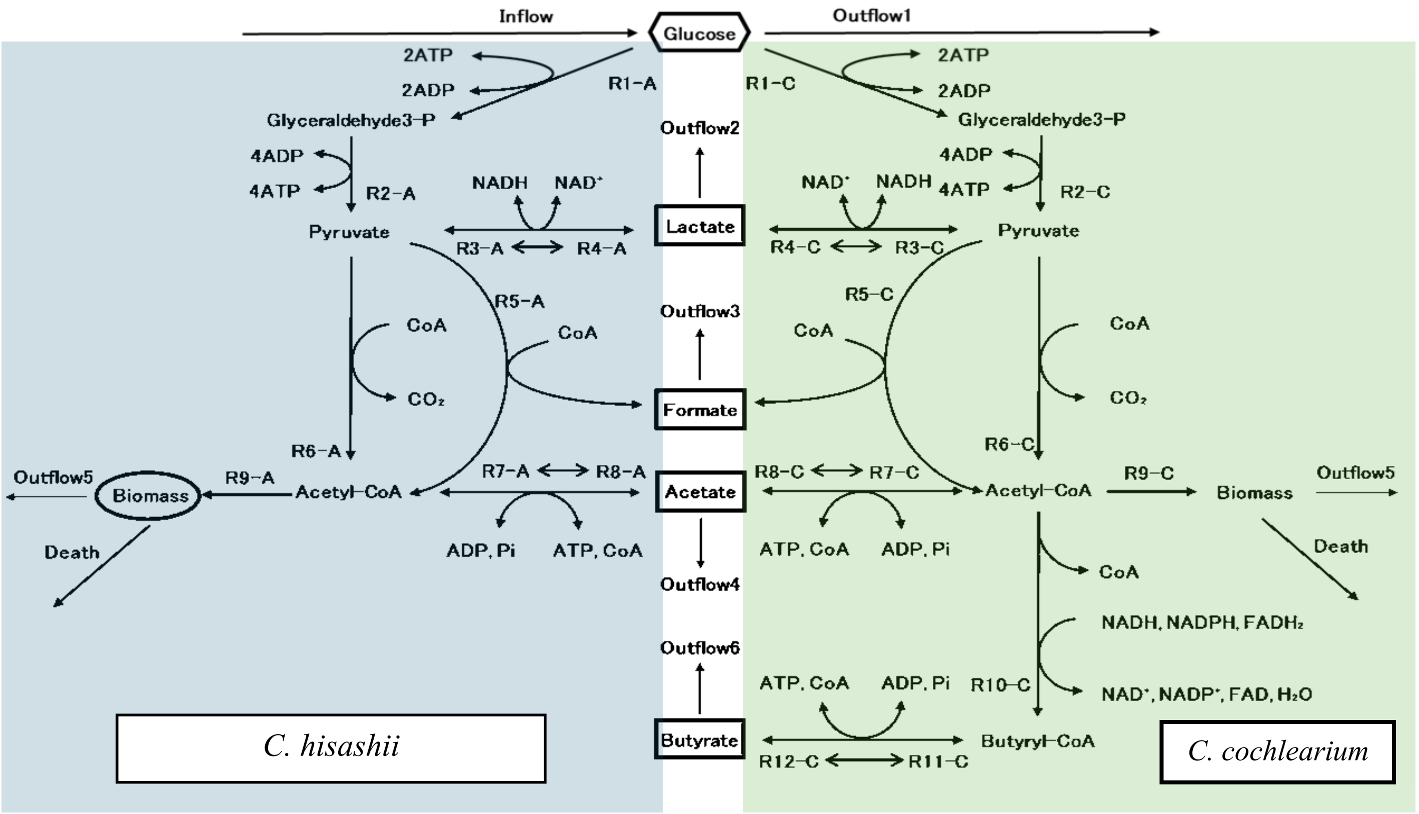


**Supplementary Fig. 1**
**Metabolic pathways of *C. hisashii* (blue) and *C. cochlearium* (green) in modified EMP pathway and organic acids productions pathway at D = 0.05 h$^{-1}$.**

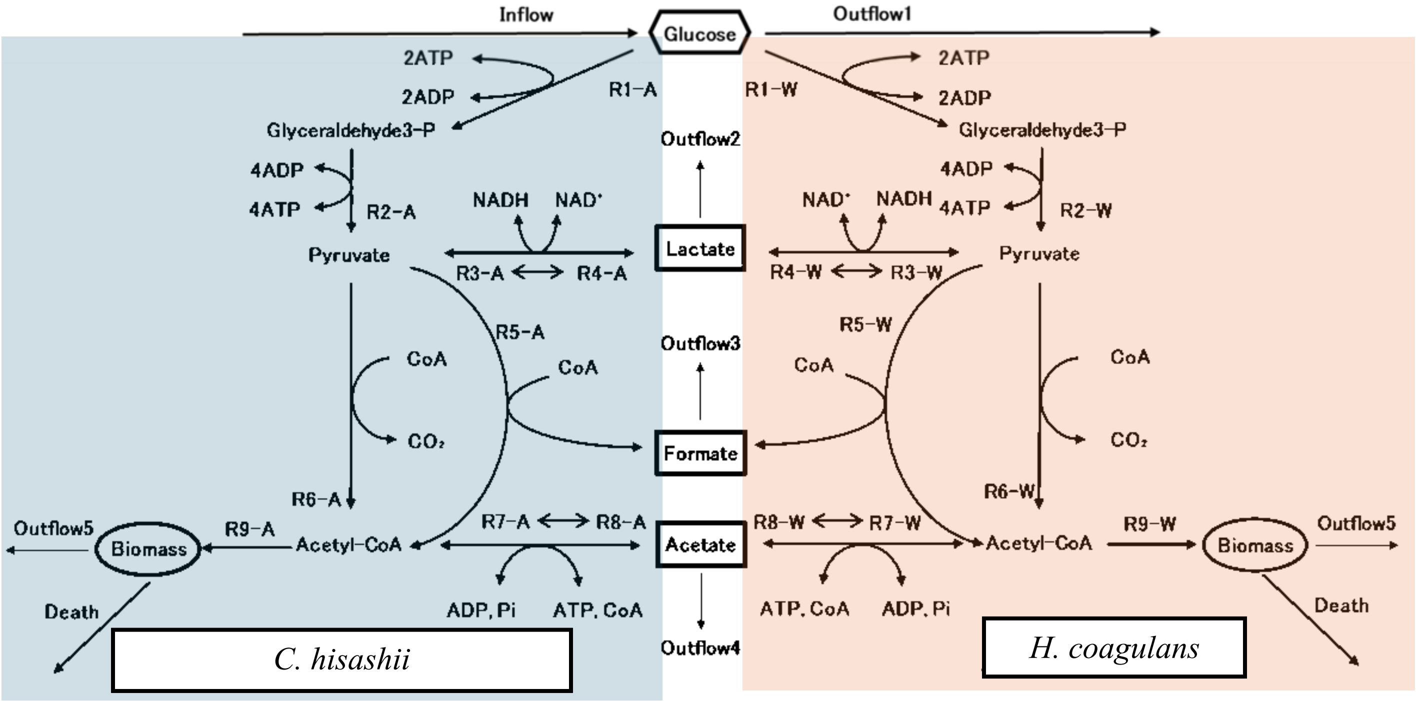


**Supplementary Fig. 2**
**Metabolic pathways of *C. hisashii* (blue) and *H. coagulans* (orange) in modified EMP pathway and organic acids productions pathway at D = 0.4 h$^{-1}$.**

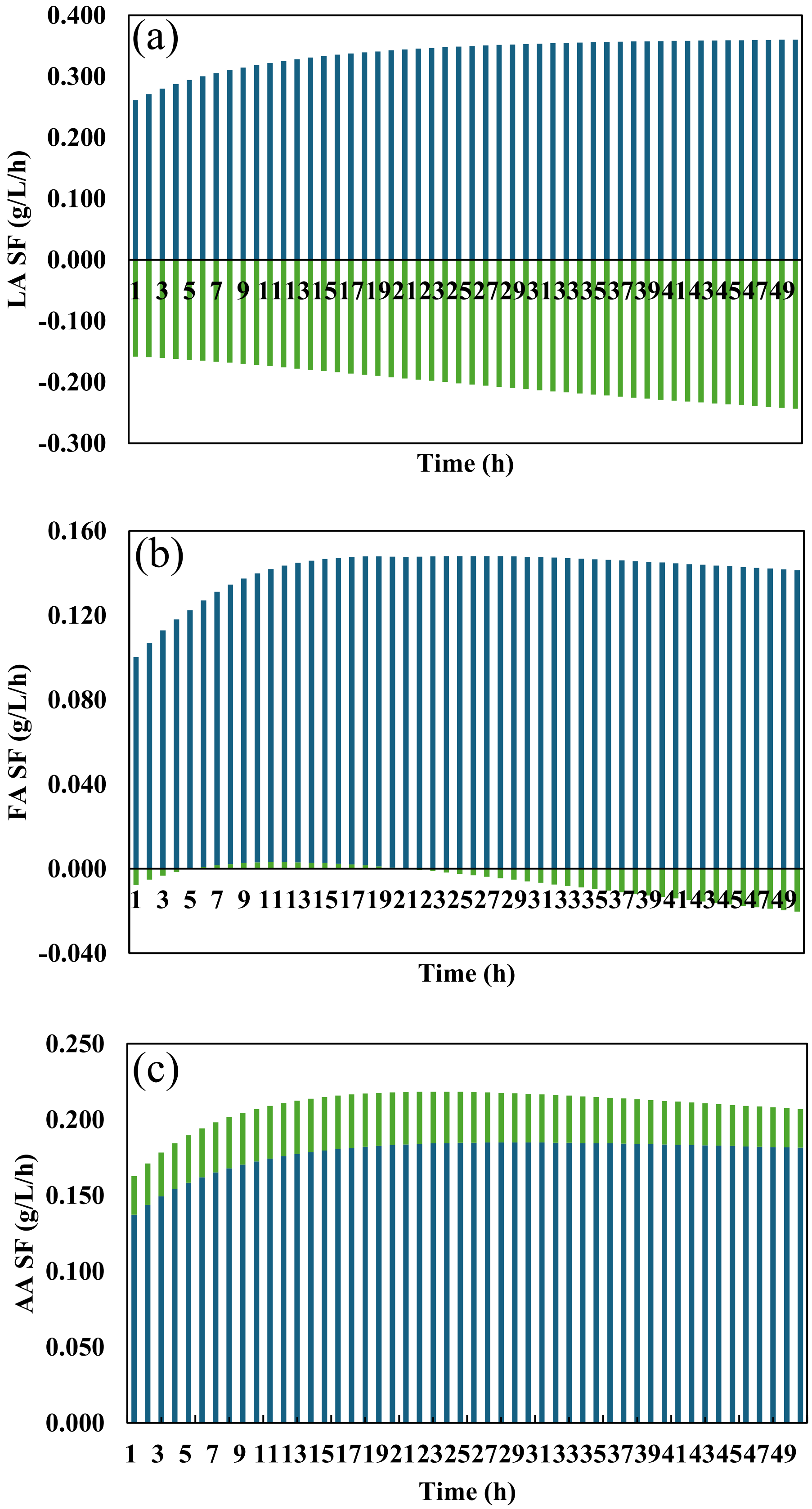


**Supplementary Fig. 3**

**Dynamics of SF in each time of (a) lactic acid, (b) formic acid, and (c) acetic acid by *C. hisashii* (blue) and *C. cochlearium* (green) at D = 0.05 $h^{-1}$.**

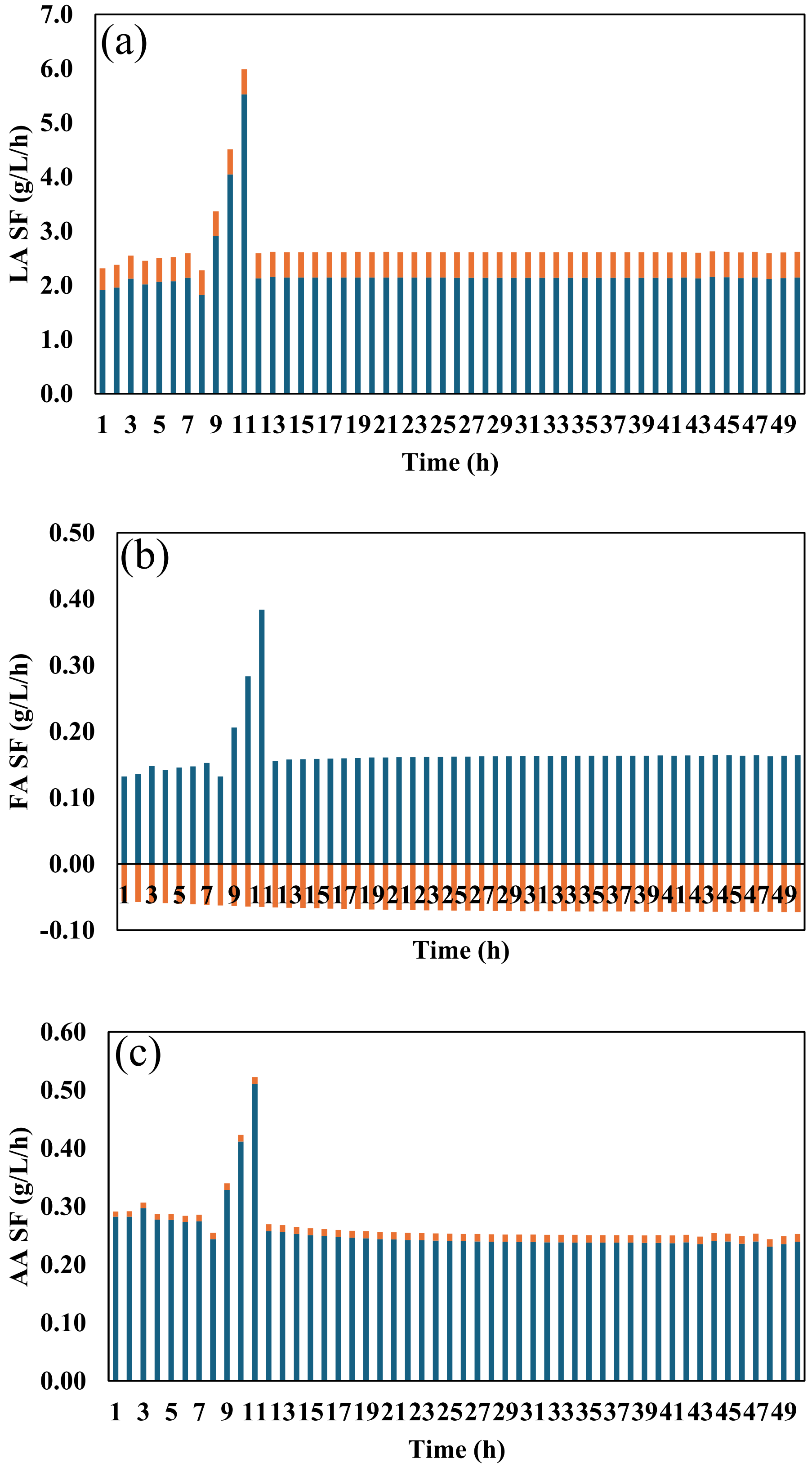


**Supplementary Fig. 4**

**Dynamics of SF in each time of (a) lactic acid, (b) formic acid, and (c) acetic acid by *C. hisashii* (blue) and *H. coagulans* (orange) at D = 0.4 h$^{-1}$.**
